# Feature Frequency Profile-based phylogenies are inaccurate

**DOI:** 10.1101/2020.06.28.176479

**Authors:** Yuanning Li, Kyle T. David, Xing-Xing Shen, Jacob L. Steenwyk, Kenneth M. Halanych, Antonis Rokas

**Affiliations:** Department of Biological Sciences, Vanderbilt University, Nashville, Tennessee, USA; Department of Biological Sciences, Auburn University, Auburn, Alabama, USA; State Key Laboratory of Rice Biology and Ministry of Agriculture Key Lab of Molecular Biology of Crop Pathogens and Insects, Institute of Insect Sciences, Zhejiang University, Hangzhou 310058, China

## Abstract

Choi and Kim (PNAS, 117: 3678-3686; first published February 4, 2020; https://doi.org/10.1073/pnas.1915766117) used the alignment-free Feature Frequency Profile (FFP) method to reconstruct a broad sketch of the tree of life based on proteome data from 4,023 taxa. The FFP-based reconstruction reports many relationships that strongly contradict the current consensus view of the tree of life and its accuracy has not been tested. Comparison of FFP with current standard approaches, such as concatenation and coalescence, using simulation analyses shows that FFP performs poorly. We conclude that the phylogeny of the tree of life reconstructed by Choi and Kim is suspect based on methodology as well as prior phylogenetic evidence.

## Main

Choi and Kim (1) used the alignment-free Feature Frequency Profile (FFP) method to reconstruct a broad sketch of the tree of life based on proteome data from 4,023 taxa. The FFP-based reconstruction reports many relationships that strongly contradict the current consensus view of the tree of life, including sister group relationships for plants + animals, Bacteria + Archaea, and Mollusca (incorrectly referred to as cnidarians) + deuterostomes. The FFP-based tree also contains unexpected placements for several “singleton” taxa, such as the position of the chordate *Ciona intestinalis* as sister to a clade including all other chordates, arthropods, mollusks, and annelids. Given that these results are based solely on the FFP method (1, 2), whose accuracy has not been tested, scrutiny is required.

The FFP method is a variation of “Word Frequency Profile”, which is commonly used in information theory and computational linguistics (3). Briefly, the FFP corresponds to a vector of the counts of unique k-mers in a DNA or amino acid sequence. To construct an FFP-based phylogenetic hypothesis, distances between different sequences are measured by Jensen-Shannon Divergence (JSD) followed by inference using BIONJ (4).

To test the performance of the FFP method, we compared it to two standard approaches of phylogenomic inference, namely maximum likelihood (ML) analyses based on concatenation and coalescence. We first measured the topological distances between trees produced by the three approaches on a 2,408-gene, 343-taxon phylogeny of budding yeasts (5). We found that the phylogenetic hypotheses inferred from concatenation and coalescence approaches shared 91.5% of bipartitions; in contrast, the phylogenetic hypothesis inferred using concatenation shared 72.4% of bipartitions with the phylogenetic hypothesis inferred using FFP, and the phylogenetic hypothesis inferred using coalescence shared 68.8% of bipartitions with the FFP hypothesis (Fig. 1A). These results suggest that FFP-based results greatly differ from those inferred by concatenation and coalescence.

**Figure 1.**
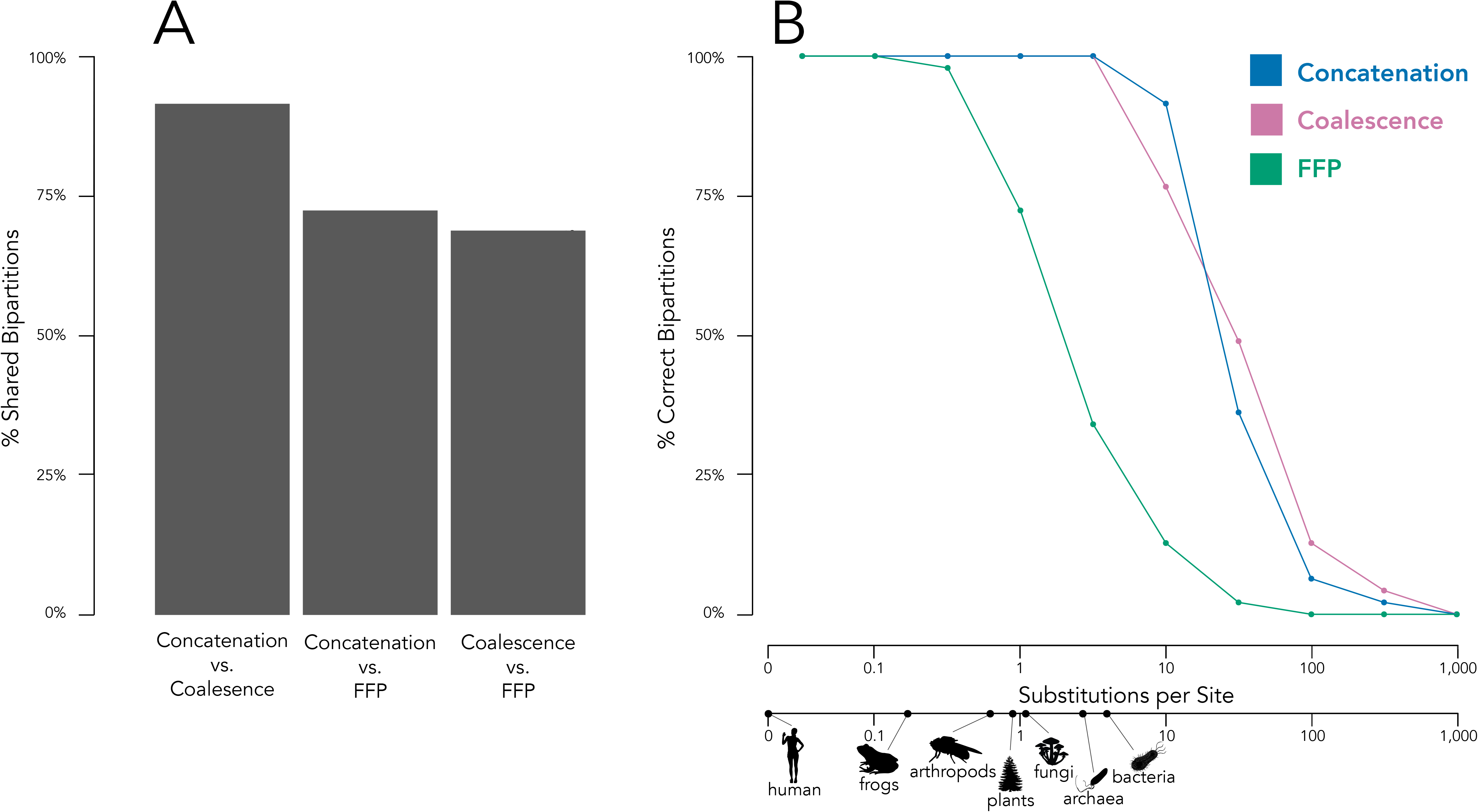
The Feature Frequency Profile (FFP) method performs poorly compared to standard approaches of statistical phylogenetic inference. (A) Topological similarities between ML-based concatenation, coalescence based on proteomes from 343 yeast taxa (5). Topological accuracy of concatenation, coalescence and FFP approaches in recovering the 50-taxon balanced tree topology used in the simulation analysis. Each data point corresponds to the average percentage of correctly inferred bipartitions from phylogenetic analyses of 100 simulated sequence alignments. The different data points represent the results of simulations using trees with different branch lengths. Silhouettes indicate the average number of amino acid substitutions/site between conserved ribosomal proteins in a reference taxon (in this case human) and other clades. Branch lengths were taken from Hug et al. (2016) (13).

To further evaluate the performance of FFP compared to standard phylogenetic methods, we simulated 100 genes under a 50-taxon balanced tree using a panel of different substitution rates and tested the accuracy of concatenation, coalescence, and FFP approaches in recovering the topology used to generate the data (Fig. 1B). We found that FFP inferred a much lower percentage of correct bipartitions than either the concatenation or coalescence approaches. FFP’s lower accuracy is particularly notable when evolutionary rates that exceed 0.5 substitutions / site are used (Fig. 1B), which are commonplace in analyses of deep phylogenies.

The discrepancy between FFP and concatenation and coalescence approaches stems from the fact that this method is not designed to infer evolutionary history (3). By measuring the overall similarity between sequences, FPP is a phenetic or similarity-based method that does not account for homoplasy stemming from the occurrence of multiple state changes over time (6, 7). Thus, it will be misled by multiple substitutions, especially over large evolutionary distances. Similarly, branch lengths in FPP trees measure similarity between sequences and should not be conflated with evolutionary distance or time.

Our analyses suggest that FFP underperforms compared to current standard approaches, such as concatenation- and coalescence-based approaches, and is a poor method for inferring the Tree of Life. As such, the phylogeny of Choi and Kim (2020) is suspect based on methodology as well as prior phylogenetic evidence.

## Supplementary Information

### Supplemental methods

All scripts and data used for our analyses will be made publicly available in a Figshare repository at 10.6084/m9.figshare.12543050 upon publication.

To examine the degree of topological similarity of phylogenies inferred with the FFP method with concatenation and coalescence phylogenies, we used the proteomes of 343 budding yeast species and outgroups from a previous study (5). Briefly, we first calculated the FFP values for each sequence for k-mer size of 13. We then measured the divergence of all pairs of FFPs using the Jensen-Shannon Divergence (JSD). The distance data matrix was used as input for neighbor joining (NJ) tree building using BIONJ with default settings (4). We quantified the degree of incongruence for every bipartition (or internal branch or internode) by considering all prevalent conflicting bipartitions among phylogenetic trees (8, 9) using the “compare” function in Gotree version 1.13.6 (https://github.com/evolbioinfo/gotree).

To evaluate the accuracy of FFP-based phylogenies relative to concatenation and coalescence phylogenies, we conducted simulation studies. All simulations used a 50-taxon balanced tree, scaled by substitution rate (Fig. 1a). Each reference tree was used to generate a data matrix with 1,000 amino acid gene alignments with 500 sites under LG model using Pyvolve v1.0.1(10).

For each simulated data matrix, we used three approaches to infer the phylogeny:

1. *a concatenation approach with a single model or partition:* For the concatenation approach, all phylogenetic analyses were performed using IQ-TREE, multi-thread version 1.6.8 (11). The topological robustness of each gene tree was evaluated by 1,000 ultrafast bootstrap replicates.
2. *a multi-species coalescent-based approach that used individual gene trees to construct the species phylogeny:* For the coalescence approach, individual gene trees were inferred using IQ-TREE with an LG model. Topological robustness of each gene tree was evaluated by 1,000 ultrafast bootstrap replicates. We used individual ML gene trees to infer the coalescent-based species tree using ASTRAL-III version 5.1.1 (12) for each data matrix. Topological robustness was evaluated using the local posterior probability (LPP).
3. *the FFP method:* For the FFP method, we calculated the FFP values for each simulated data matrix, and the JSD and NJ tree was conducted using the same settings as above.

For all the phylogenies inferred from the simulation data matrices, the degree of topological accuracy (i.e., the degree of topological similarity to the reference tree used to simulate the sequence alignments) was quantified by measuring the degree of incongruence for every bipartition (or internal branch or internode) by comparing all prevalent conflicting bipartitions between the reference tree and the inferred tree (8, 9) using the “compare” function in Gotree version 1.13.6 (https://github.com/evolbioinfo/gotree).

## Notes

### Competing Interest Statement

The authors have declared no competing interest.

